# Spontaneous thought and microstate activity modulation by social imitation

**DOI:** 10.1101/2021.01.15.426876

**Authors:** Miralena I. Tomescu, Claudiu C. Papasteri, Alexandra Sofonea, Romina Boldasu, Valeria Kebets, Catalina Poalelungi, Ioana R. Podina, Catalin I. Nedelcea, Alexandru I. Berceanu, Ioana Carcea

## Abstract

Social imitation increases well-being and closeness by mechanisms that remain poorly understood. We propose that imitation impacts behavioural states in part by modulating post-imitation mind-wandering. The human mind wanders spontaneously and frequently, revisiting the past and imagining the future of self and of others. External and internal factors can influence wandering spontaneous thoughts, whose content predicts subsequent emotional states. In 43 young subjects, we find that imitating the arm movements of an actor alters the dynamics and the content of subsequent resting-state spontaneous thoughts. Imitation-sensitive features of spontaneous thoughts correlate with both behavioural states and salivary oxytocin levels. EEG microstate analysis reveals that global patterns of correlated neuronal activity predict imitation-induced changes in spontaneous thoughts. Thus, imitation can modulate ongoing activity in specific neural networks to change spontaneous thought patterns as a function of oxytocin levels, and to ultimately orchestrate behavioural states.

## Introduction

Humans spend a significant fraction of their waking time defaulting to ‘mind-wandering’, an unconstrained succession of mental states that generate spontaneous thoughts^1–4^. The content of spontaneous thought varies widely within and between individuals, and can be characterized with respect to its dynamics (more or less fragmented), affect (negative, neutral, positive), temporal orientation (remembering or planning), social orientation (self or others), mental modality (visual or verbal), and association with physiological states (sleepiness, stress, etc)^3, 5–8^. Many of these dimensions can be captured in experimental settings by validated retrospective self-reported questionnaires, like the Amsterdam Resting-State Questionnaire (ARSQ)^9^.

Spontaneous thoughts correlate with explicit affective state. Negative mood and spontaneous thoughts interact circularly, as sad spontaneous thoughts tend to be both preceded and succeeded by sad or anxious moods^7,10^. Socio-temporal features of spontaneous thoughts can also predict subsequent mood, where past- and other-oriented thoughts predict negative affect, but future- and self-oriented thoughts predict improved mood^7^. Other studies, however, found that self-related content of spontaneous thoughts correlates with sad mood and with depressive symptoms^11^.

Spontaneous thoughts can serve important roles in mental well-being, including in generating personal goals^12–14^, in memory consolidation^15,16^, and in fostering creativity^14, 16–18^. In other words, spontaneous thought profiles can improve behavioural states and serve as internal tools for cognitive and emotional well-being. Could these tools be leveraged in behavioural interventions? This possibility is bolstered by the documented impact of contextual factors on spontaneous thoughts^8^. However, much more remains to be understood about the nature of contexts capable of changing mind-wandering, and about the mechanisms by which they achieve such changes.

Spontaneous thoughts likely arise from task-independent ongoing brain activity^19,20^. Functional imaging implicates several brain networks in spontaneous thought. Primarily, the default mode network (DMN), which is active during states of rest, and decreases as a consequence of task demands^20–22^ is closely linked to the generation of spontaneous thoughts^23^. In addition to the DMN, mind-wandering also engages salience and executive control networks^21,24,25^. These networks are believed to impose automatic and deliberate constraints on mind-wandering, or in other words they track the ‘wandering path’ from one spontaneous thought to another^4^.

The logistics of functional imaging (laying still in a loud scanner) might bias the nature of spontaneous thoughts. A more naturalistic approach to probing the neural substrates of mind-wandering is to perform scalp electroencephalographic (EEG) recordings, and characterize activity-correlated networks using microstate analysis. This analysis identifies transiently (60-120 ms) quasi-stable global patterns of scalp potential topographies that are highly reproducible within and across subjects^26,27^. Between four and seven such states have been identified so far, and have been linked to fMRI resting-state networks^26,28,29,30,31^. Several studies using simultaneous EEG and fMRI, or direct EEG source localization methodology, support the notion that microstates A and B are associated with primary sensory brain regions like visual, auditory/language cortices, while C, D, and E microstates associate with core regions of the posterior DMN, attention/cognitive control, and salience resting-state networks, respectively ^26,31^. Potential roles of these microstates in visual imagery, cognitive control and planning have been described and also disputed^26^. However, a distinct pattern of modulations between the posterior DMN C state and attention D state might reinforce their functional meaning. Relaxed, meditative, and hypnotic states entrain longer and more frequent posterior DMN C states, and slower attention D temporal dynamics^26,32–35^. Microstate analysis could deepen the understanding of neural substrates for spontaneous thoughts. It could also identify mechanisms by which contextual factors change the pattern and spontaneous thoughts and their effects on behaviour.

Social factors represent a major source of context variation in humans, however little is known about the relationship between social contexts and spontaneous thoughts. In the current study, we hypothesize that social contexts can change subsequent resting-state spontaneous thoughts, and microstate activity patterns. To test this, we used a dyadic social imitation task, where subjects follow the arm movements of an actor. We previously showed that this form of social imitation decreases momentary stress, increases well-being and social closeness^36^. Several mechanisms for these behavioural changes have been proposed, including rewarding effects, and perception-action matching ^36,37^. Hormonal substrates have also been implicated, particularly increased levels of oxytocin^36,38^. We reasoned that in addition to the above mechanisms, imitation could modify behavioural states indirectly, by changing ongoing brain activity and the pattern of spontaneous thoughts. We used a combination of behavioural, biochemical and physiological measures to show that spontaneous thought patterns and ongoing activity of neural networks are sensitive to social imitation. Our findings indicate a potential mechanism by which social imitation, which is already used in theatre as an exercise that decreases stage fright and increases closeness between actors, could be therapeutic in the general population.

## Results

To determine if brief dyadic social imitation (SI) might alter spontaneous thoughts and ongoing activity of neural network in imitators, we probed five-minute resting-state episodes before (PRE) and after (POST) the task (**Fig. 1a**). The SI task consisted of a three-minute interaction where the subject followed the arm movements of an actor, as previously described^36^. For the control, non-social condition (CTRL), subjects were asked to depict with their arm movements the shape of geometrical figures displayed on a computer screen (**Fig. 1b**). Within each subject, experimental and control conditions were completed on separate days, and in random order.

**Figure 1.**
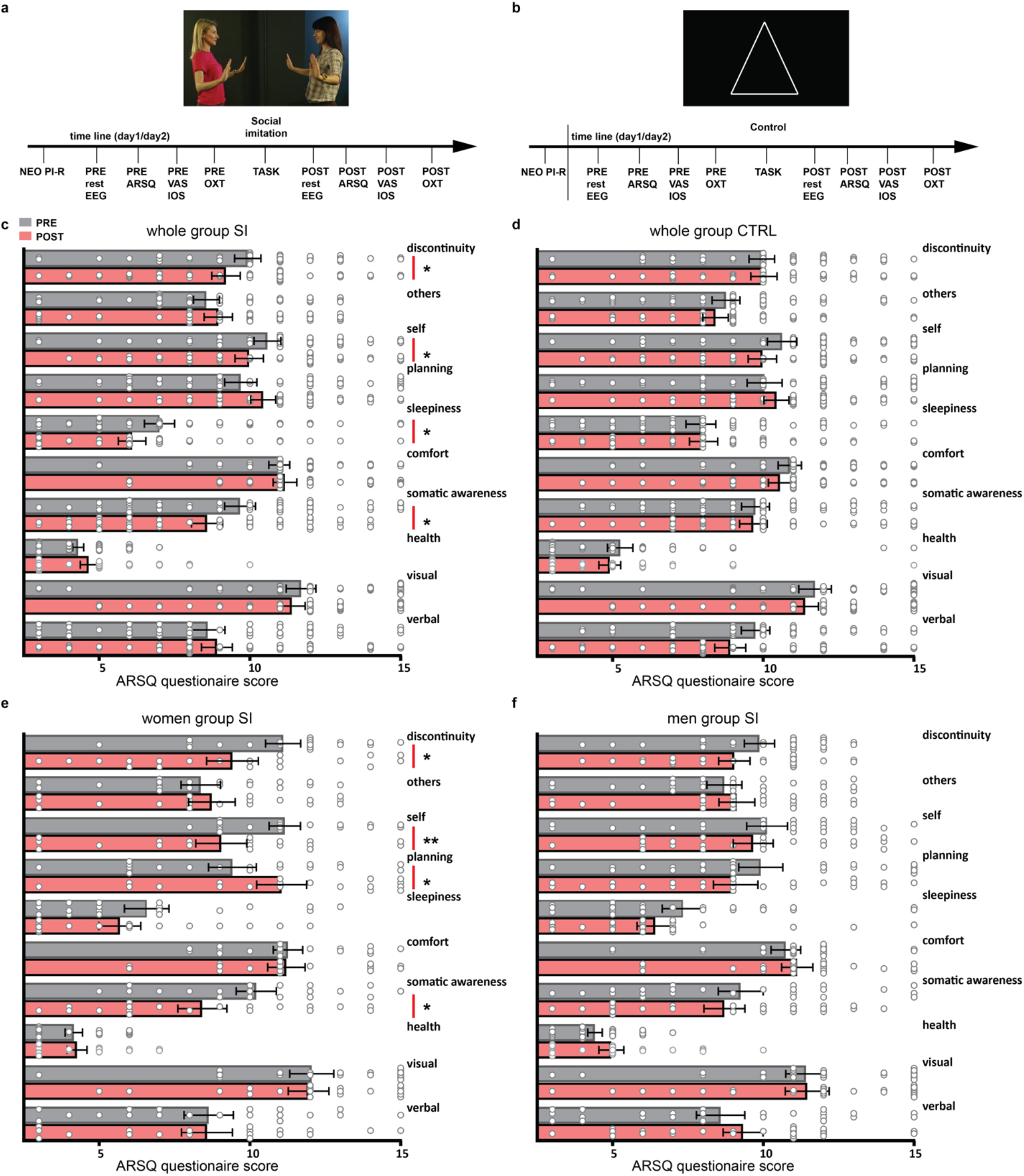
SI changes spontaneous thoughts. (**a**) Experimental design, SI task. (**b**) Design of control task. (**c**) Resting-state ARSQ scores before (PRE, gray) and after (POST, red) the SI task. (**d**) Restingstate ARSQ scores before and after CTRL task. (**e**) Effects of SI on ARSQ scores in women. (**f**) Effects of SI on ARSQ scores in men. Errors are SEM, *p<0.05, **p<0.001.

### Modulation of resting-state spontaneous thoughts by social imitation

To quantify the changes in spontaneous-thoughts during resting-state episodes, we evaluated intrinsic mentation using a standardized test, the ARSQ 2.0 questionnaire^9^ that allowed us to construct a repertoire of the individual subjective mind-wandering experience and their modulations by tasks. In order to investigate if and how SI modulates spontaneous thoughts, we performed three-way repeated measures ANOVA with the ARSQ factors, by time (PRE vs POST) and condition (SI vs CTRL). At the group level, as expected from the structure of the test, we found a main effect of ARSQ factors (F(3, 378)=38.2, p<0.00001, 0.47 η_p_^2^). We also found a significant main effect of time (F(1,42)=4.38, p=0.042, 0.09 η_p_^2^, PRE=9.30±0.17, POST=9.05 ±0.18, N=43), and a significant three way interaction between time, ARSQ factors and conditions (F(9,378)=2.03, p=0.03, 0.04 η_p_^2^). After the SI task, subjects reported less fragmented thoughts, fewer thoughts about themselves and their bodies, and felt less tired (**Fig. 1c**). Post-hoc tests revealed significant changes induced by SI in four different factor categories: *discontinuity of mind* (PRE= 10.41±2.53, POST=9.20±3.09, p=0.005, N=43), *self-related thoughts* (PRE=10.58±2.91, POST=9.39±3.4, p=0.0006), *sleepiness* (PRE=7.0±3.28, POST=6.09±2.9, p=0.036) and *somatic awareness* (PRE=9.67±3.34, POST=8.58±3.38, p=0.012) (**Fig. 1c**). No significant changes were detected after the CTRL conditions (**Fig. 1d**). This indicates that imitating another person, but not an inanimate screen, can lead to changes in subsequent spontaneous thought dynamics (less fragmented), content (less about self), and association with physiological states (less tired).

When we separated the group by gender, we found that women account for most of the SI-induced changes in spontaneous thoughts. There was a significant main effect of time PRE-POST (F(1,18)=4.7, p=0.043, 0.20 η_p_^2^, PRE=9.45±0.28, POST=9.11 ±0.29), and a significant three way interaction between time, ARSQ factors and conditions (F(9,162)=2.83, p=0.03, 0.04 η_p_^2^). Following SI, women reported having significantly less fragmented thoughts (*discontinuity*, PRE= 11.1±2.53, POST=9.42±3.7, p=0.004), thinking less about themselves (*self*, PRE=11.1±2.26 POST=9.05±3.6, p=0.0003) and their bodies (*somatic awareness*, PRE=10.21.67±2.89, POST=8.42±3.54, p=0.002) (**Fig. 1e**). Moreover, women reported a significant increase of the *planning* factor (PRE=9.42±3.45, POST=11.06, p=0.005) (**Fig. 1e**). On the contrary, in men we found neither a main effect of time (F(1,22)=0.6, p=0.44) nor a significant interaction between time, ARSQ factors and conditions (F(9,198)=1.48, p=1.56) (**Fig. 1f**).

### Associations between spontaneous thoughts and behavioural states

We previously showed that SI but not the CTRL task can increase well-being and social closeness, whereas both SI and CTRL conditions lead to decreased self-reported stress^36^. To determine if there is an association between these momentary self-reported behavioural states and spontaneous thoughts, we performed rank correlation analyses between behavioural and ARSQ scores across all conditions (**Fig. 2**). In women we found a significant positive correlation between *discontinuity of mind* and perceived stress level (Gamma r=0.36, p=0.033), that was not significant either at the group level or in men (**Fig. 2a**). We also found a positive association between the level of perceived stress and *self*-oriented thoughts (Gamma r=0.35, p=0.001, **Fig. 2b**) and thoughts about *health* (Gamma r=0.29, p=0.007, **Fig. 2d**). Stress was negatively associated with thoughts about *comfort* (Gamma r= −0.31, p=0.004, **Fig. 2d**). The reversed pattern of correlation was found for wellbeing (*health*: Gamma r= −0.33, p= 0.002; *comfort*: Gamma r=0.56, p<0.0001, **Fig 2c,d**). Thoughts about future *plans* were positively associated with both closeness (Gamma r=0.24, p=0.02) and well-being (Gamma r=0.21, p= 0.04) (**Fig. 2d**). These findings indicate a relationship between behavioural states and the pattern of spontaneous thoughts, both of which change after SI, particularly in women.

**Figure 2.**
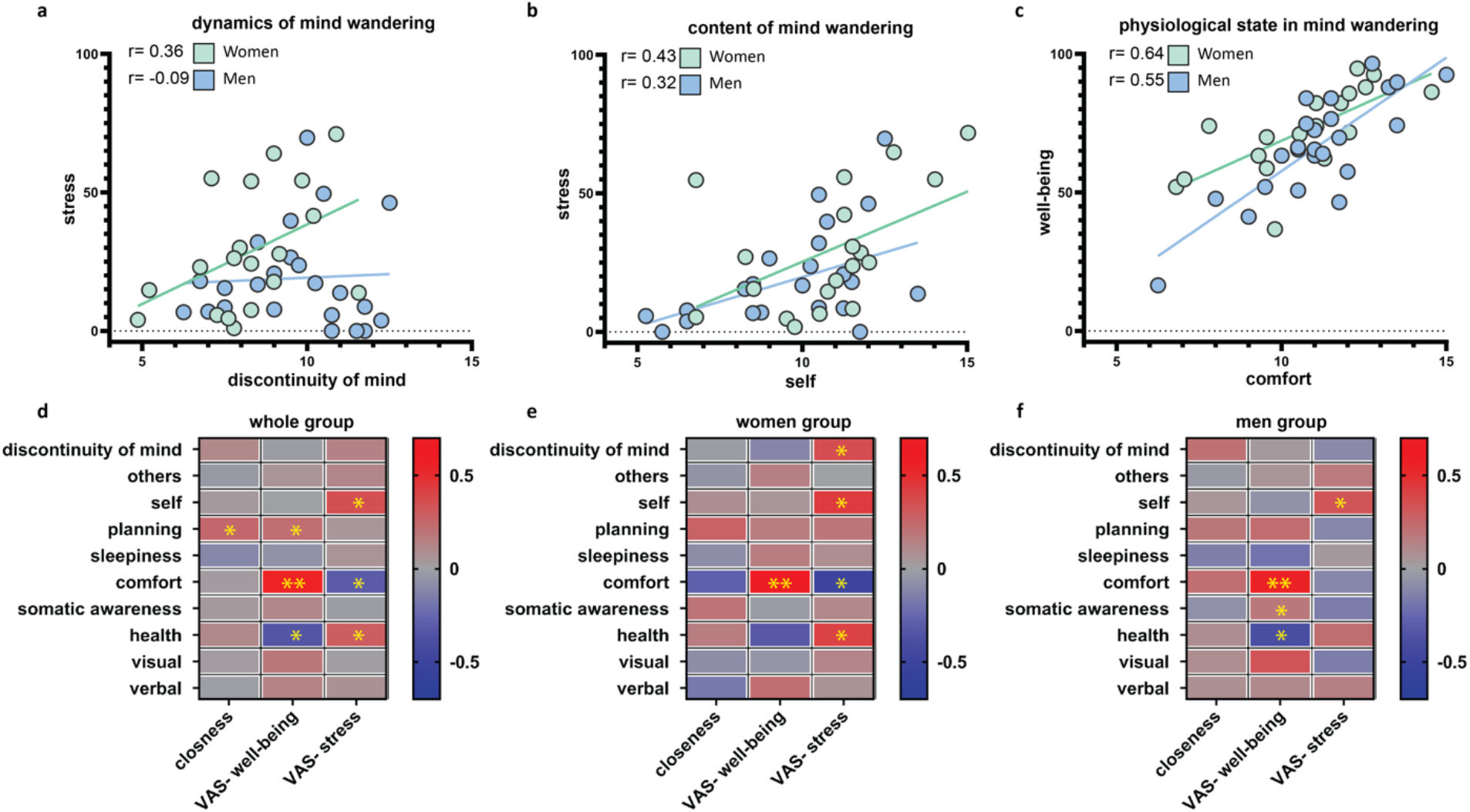
The pattern of spontaneous thoughts associates with subjective behavioral states. (**a**) Dynamics of spontaneous thought positively associate with self-reported stress levels. (**b**) Self-centered thought content positively correlates with stress. (**c**) Perceived states of ‘comfort’ during mind-wandering positively correlate with self-reported well-being. (**d**) Summary data of correlations (r values) between ARSQ scores and self-reported behavioral state measures for the entire group. (**e**) Summary of correlations for women. (**f**) Summary of correlations for men, *p<0.05, **p<0.001.

### Associations between spontaneous thoughts and salivary oxytocin

Imitation has been linked to increased levels of salivary oxytocin^36,38^. We previously showed that this increase is only partially related to the outcome of the SI on self-reported behavioural states^36^. In order to find how spontaneous thoughts across conditions relate to salivary oxytocin levels, we calculated correlations that showed several significant negative associations of salivary OXT dynamics with *discontinuity of mind* (Gamma, r=-0.25, p= 0.029, **Fig. 3a**), with thoughts about future *planning* (Gamma, r=-0.25, p= 0.03, **Fig. 3b**) and, only in the men group, with *comfort* (Gamma, r=-0.25, p= 0.03) (**Fig. 3c**). Therefore, while increased salivary oxytocin could represent a potential mechanism for changing the dynamics of spontaneous thoughts (i.e., less fragmented thinking), it does not appear to affect their content or their association with physiological states.

**Figure 3.**
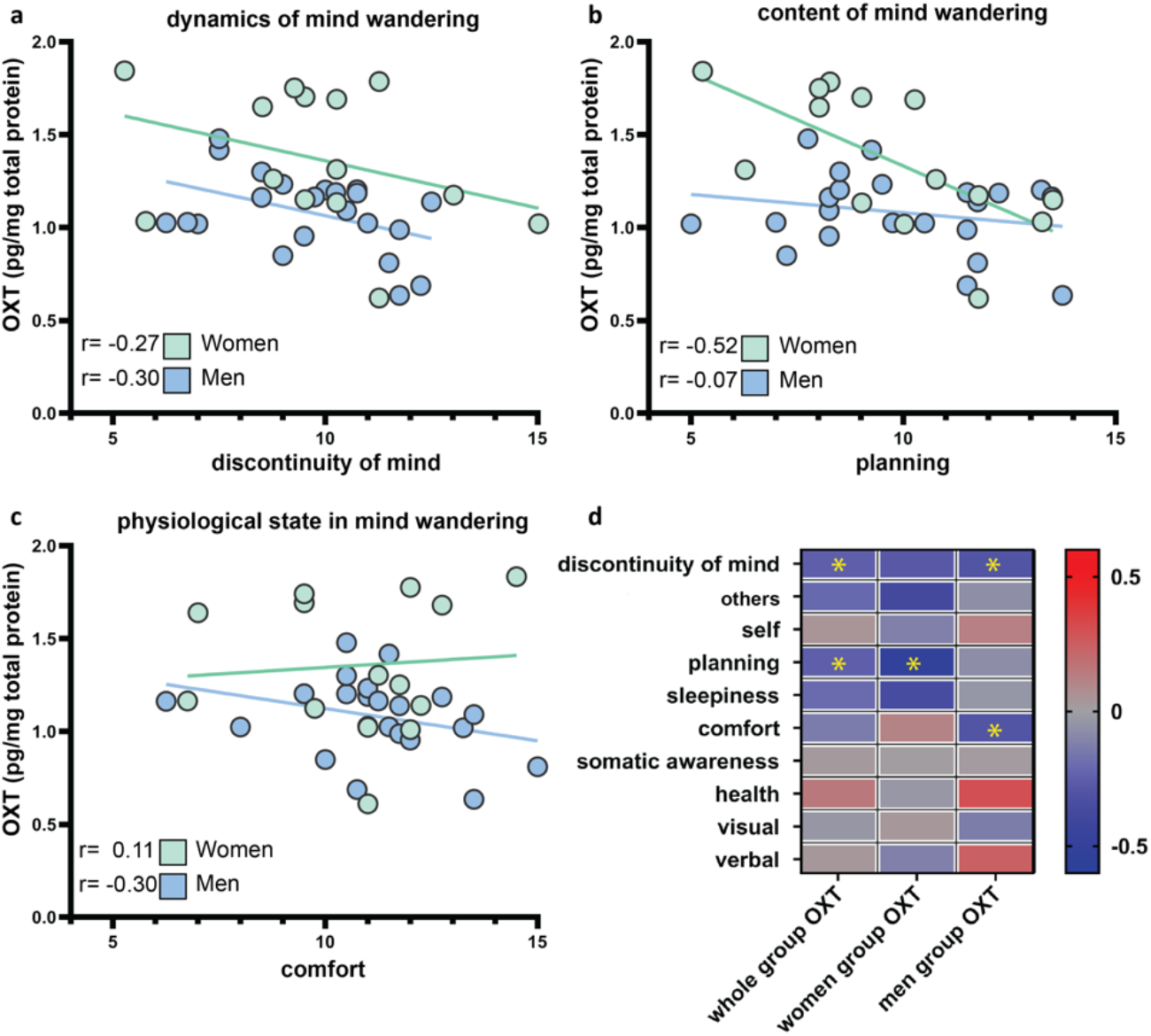
Spontaneous thought patterns associate with salivary OXT levels. (**a**) Dynamics of mind-wandering negatively correlate with salivary OXT. (**b**) Thoughts about ‘planning’ negatively correlate with OXT in women. (**c**) States of ‘comfort’ negatively correlate with OXT. (**d**) Summary data of correlations between ARSQ scores and salivary OXT, *p<0.05.

### Neural correlates for SI-induced changes in resting-state activity

To investigate the neural substrate for SI-induced changes in mentation, we recorded EEG activity during resting-state. Using a set of seven different independent criteria (described in **Methods**), we determined that five microstates can optimally describe group topographical variability. Across all conditions (PRE, POST before and after the SI/CTRL) and individuals (N=43), summing more than 800 individual resting-state dominant topographies, the cluster analysis robustly identified A, B, C, D and E prototypical microstates that explained 81.9 % of variance (**Fig. 4a**). We then analysed SI-induced changes in the duration and occurrence of these EEG microstates.

**Figure 4.**
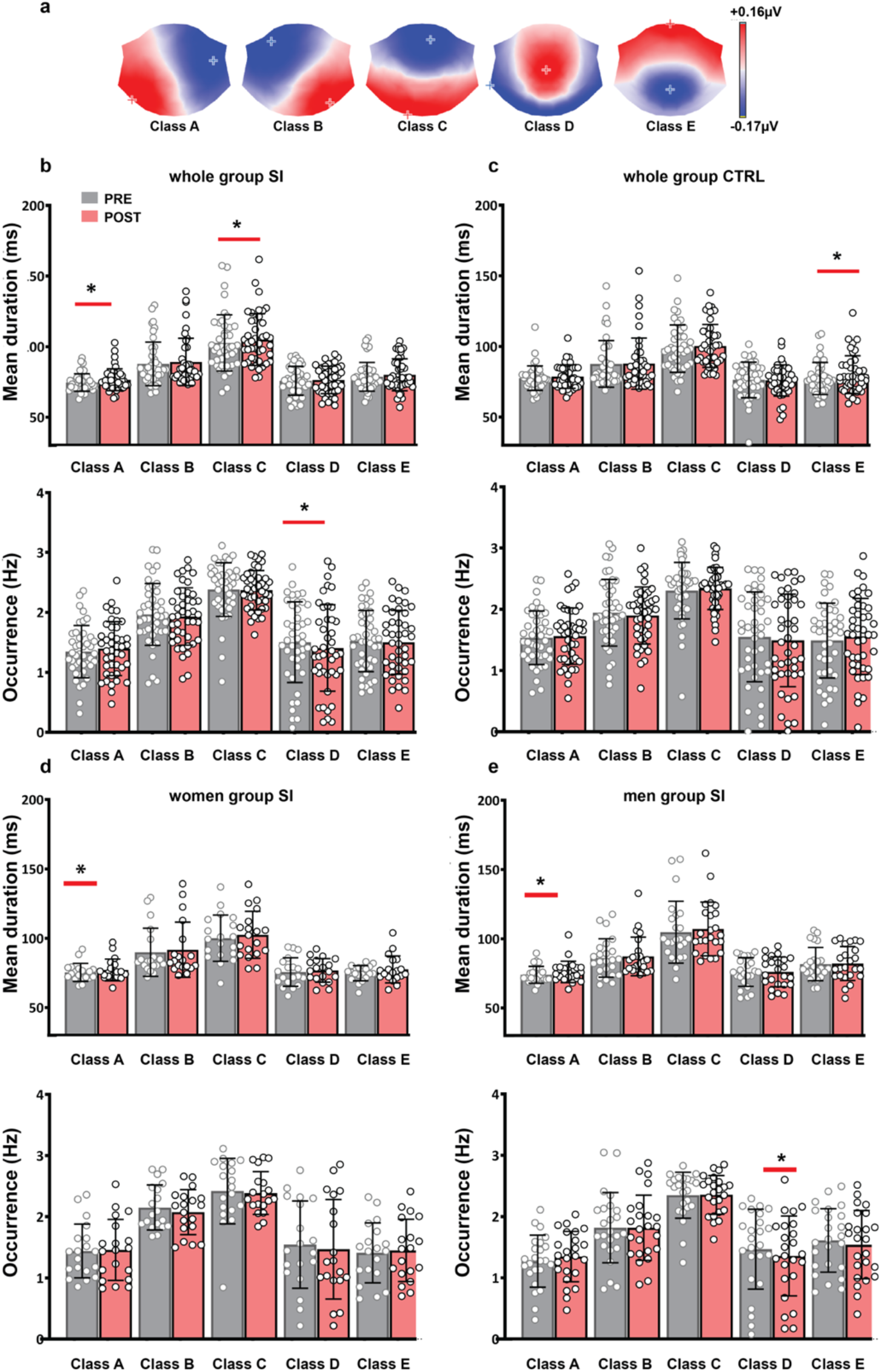
Microstate dynamics following social imitation. (**a**) Identified classes of microstates. (**b**) SI-induced changes in mean duration (top) and occurrence (bottom) for each class. (**c**) Changes in mean duration (top) and occurrence (bottom) following CTRL task. (**d**) SI-induced changes in class duration (top) and occurrence (bottom) in women. (**e**) SI-induced changes in class duration (top) and occurrence (bottom) in men, *p<0.05.

### SI modulates the intrinsic temporal dynamics of microstates

To identify how the SI task might modulate intrinsic patterns of resting networks dynamics, we performed repeated-measures three-way ANOVA analysis on the *duration* (*ms*) and *occurrence* (*Hz*) of microstates, by time (PRE vs POST) and by condition (SI vs CTRL). Separate paired Wilcoxon tests were performed to identify the specific task modulation for each microstate.

In the ANOVA analysis for microstate *duration* we found a main effect of time (N=43, PRE=83.8±5.50, POST=85.45±5.55, F(1,42)=10.7, p=0.002, 0.20 η_p_^2^) and a main effect of microstates (A=75.6.±6.71, B=88.5±16.06, C=103.71±18.71, D=76.19±9.84, E=79.16±9.84, F(1,42)=36.53, p<0.00001, 0.46 η_p_^2^, **Fig. 4b**). After the SI task, two classes of microstates showed increased *duration*, microstate A (PRE=74.5±6.5, POST=76.5±8.1, p=0.006) and microstate *C* (PRE=102.6, ±20.3, POST=105±18.7, p=0.04, **Fig. 4b**). We also observed a significant increase in microstate E *duration* after the CTRL task (PRE=77.3±11.3, POST=80±13.3, p=0.006, **Fig. 4c**). Thus, as hypothesized, the SI task modulated a different pattern of network dynamics than the CTRL task.

Regarding *occurrence*, we observe neither a significant ANOVA time effect nor a significant time, condition and microstate interaction. However, we found a main effect of microstates (A=1.38.±0.41, B=1.95±0.48, C=2.37±0.35, D=1.46±0.69, E=1.49±0.48 F(4,168)=30.7, p<0.00001, 0.42 η_p_^2^), and a significant time by microstate interaction (A-PRE=1.44±0.37 vs POST=1.49±0.39, B-PRE=1.95±0.47 vs POST=1.91±0.43, C – PRE=2.33±0.35 vs POST=2.35±0.27, D-PRE=1.53±0.62 vs POST=1.45±0.67, E –PRE-1.49±0.49 vs POST=1.52±0.5, F(4,168)=2.47, p=0.04, 0.05η_p_^2^). Post-hoc tests showed that SI specifically decreases the occurrence of microstate D (PRE=1.51 ±0.67, POST=1.41±0.72, p=0.012, **Fig. 4b**).

In women, we found a substantial effect of time (N=19, PRE=82.12±4.11, POST=85.12±5.10, F (1,18)=11.9, p=0.002, 0.39 η_p_^2^), a main effect of condition (SI= 84.12±4.42, CTRL=82.40±4.21, F=(1,18)=9.54, p=0.006, 0.34 η_p_^2^) and a main effect of microstates *duration* (A=76.24.±6.94, B=90.78±18.5, C=101.24±16.17, D=76.2±9.2, E=76.15±7.3 F(4, 72)=12.57, p<0.0001, 0.41 η_p_^2^). At the level of paired Wilcoxon analyses, we found a significant POST SI increased *duration* of microstates A (N=19, PRE=75.36±6.47, POST=77.13±7.8, p=0.04), and a trend for increased E microstates *duration* (N=19, PRE=74.77±5.48, POST=77.52±9.73, p=0.07, **Fig. 4d)**. After the CTRL task we found a significant increased E microstates *duration* (N=19, PRE=73.95±9.15, POST=76.28±9.36, p=0.02).

In men, we found a significant main effect of microstates (N=24, A=75.01.±6.51, B=86.67±13.59, C=105.9±20.26, D=76±10.36, E=81.89±10.86, F(4,88)=31.12, p<0.00001). At the level of paired Wilcoxon analyses, there was a significant difference between PRE and POST SI microstates class A *duration* in men (N=24, PRE=73.96±6, POST=76.05±7.73, p=0.04), and also a trend for class C increased duration (N=24, PRE=104.72±22.3, POST=107±19.44 p=0.063, **Fig. 4e**). After the CTRL task we found a trend for increased *E microstates duration* (N=24, PRE=79.95±12.42, POST=83.01±15.31, p=0.09).

With respect to microstate *occurrence* for women and men, we found a main effect of microstates in both women (A=1.53.±0.39, B=2.10±0.39, C=2.34±0.33, D=1.48±0.76, E=1.43±0.49 F(4,72)=12.1, p<0.00001, 0.40 η_p_^2^, **Fig. 4d**) and men (A=1.39.±0.33, B=1.79±0.43, C=2.34±0.25, D=1.48±0.52, E=1.57±0.46 F(4,84)=24.5, p<0.0001, 0.53η_p_^2^, **Fig. 4e**). The separate paired Wilcoxon analyses by gender showed decreased occurrence for class D only in men (PRE=1.46±0.65, POST=1.35±0.65, p=0.005, **Fig. 4e**).

### Microstate transition probabilities

In addition to the temporal characteristics for each microstate, we also investigated if SI affects the probability of transitions between microstates. For each subject we computed the number of transitions from each of the five classes to any of the other classes. These values were then normalized to all between-class transitions to obtain the final fractions for each of the twenty transitions pair (A→B, A→C, A→D, A→E, etc). The repeated three-way ANOVA revealed a main effect of time (PRE=0.0482±0.0012 vs POST=0.0486±0.0009 F(1,42)=10.9, p=0.001, 0.20 η_p_^2^), and a main effect of transition pair F(19,798)= 25.9, p<0.00001, 0.38 η_p_^2^). In addition, we identified several two-way significant interactions: the time by transition pair (F(19,798)=0.001, 0.05 η_p_^2^), and the condition by transition (F(19, 798)=1.63, p=0.04, 0.03 η_p_^2^). The B→A (SI=0.045±0.02, CTRL=0.053±0.03, p=0.048), B→C (SI=0.092±0.04, CTRL=0.081±0.04, p=0.01) and C→B transitions (SI=0.091±0.04, CTRL=0.082±0.04, p=0.02) differed significantly between the SI and CTRL conditions. Whereas the A→C (PRE=0.052±0.01, POST=0.056±0.02, p=0.021, C→A (PRE=0.052±0.01, POST=0.056±0.01, p=0.021), C→D (PRE=0.059±0.02, POST=0.056±0.02, p=0.031), C→E (PRE=0.062±0.03, POST=0.067±0.03, p=0.0008) and E→C (PRE=0.063±0.03, POST=0.067±0.03, p=0.0062) differed significantly between PRE and POST.

Separate paired Wilcoxon analyses enabled the identification of transition pairs that showed significant modulations by the SI task (**Fig. 5a-d**). There were several microstate pairs that showed increased transition after the SI: A→B (PRE=0.043±0.02, POST=0.046±0.02, p=0.019), C→E (PRE=0.063±0.04, POST=0.069±0.04, p=0.034) and E→C (PRE=0.067±0.04, POST=0.07±0.03, p=0.034). We also noted decreased frequency of transitions between C and D microstates: C→D (PRE=0.058±0.02, POST=0.056±0.02, p=0.04), D→C (PRE=0.058±0.02, POST=0.055±0.02, p=0.031) (**Fig. 5a**). However, we also found significant transition modulations following the CTRL task, when C→D decreased (PRE=0.061±0.03, POST=0.057±0.03, p=0.023), and C→E (PRE=0.061±0.03, POST=0.066±0.03, p=0.0063) and E→C (PRE=0.058±0.03, POST=0.065±0.03, p=0.027) increased in frequency (**Fig. 5b**).

**Figure 5.**
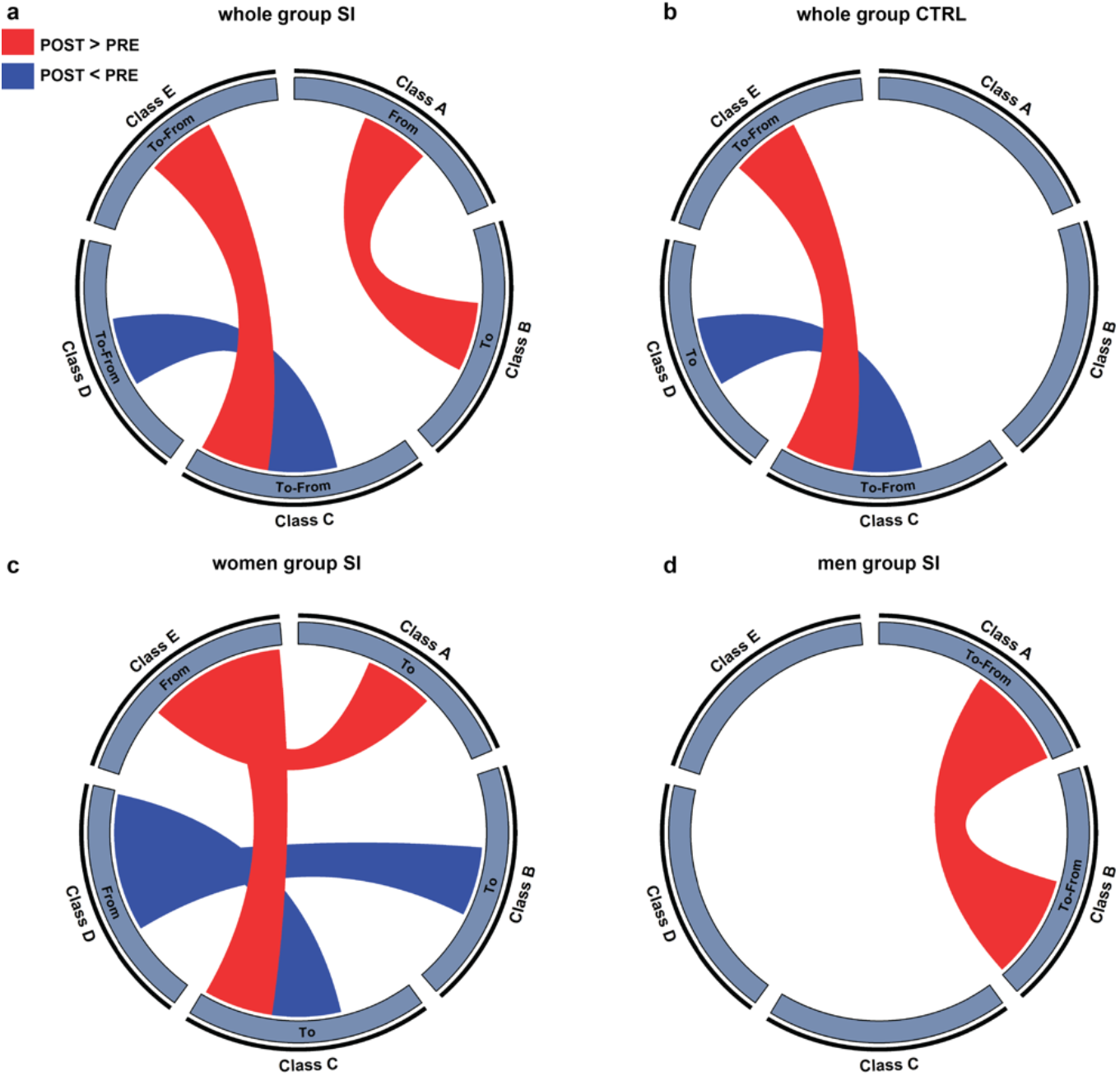
Microstate transition probability following social imitation. (**a**) Significant changes in transition probability following SI task. (**b**) Significant changes in transition probability following CTRL task. (**c**) Significant changes in transition probability following SI in women. (**d**) Significant changes in transition probability following SI in men.

In gender subgroups we found a main effect of transition pair in both women (F(19, 342)=11.28, p<0.00001, 0.38 η_p_^2^) and men (F(19, 437)=20.62, p<0.00001, 0.47 η_p_^2^), and a main effect of time in both women (PRE=0.048±0.001, POST=0.044±0.001, F(1,18)=6.97, p=0.01, 0.27 η_p_^2^) and men (PRE=0.048±0.001, POST=0.044±0.001 – F (1,23)= 5.28, 0.031, 0.18 η_p_^2^). Additionally, in men we found a main effect of condition (SI=0.0483±0.001, CTRL=0.0488±0.0007, F(1,23)=4.30, p=0.049, 0.15 η_p_^2^), and a significant two way (time by transition) interaction (F(1,437)=1.67, p=0.037, 0.06 η_p_^2^, A→C PRE=0.051±0.01, POST=0.056±0.01, p=0.02, C→E PRE=0.069±0.03, POST=0.037±0.01, p=0.01). Finally, in the men group we also found a three-way interaction, for time by condition by microstate transition (F(19, 437)=1.62, p=0.04, 0.06 η_p_^2^, **Fig. 5c,d**).

### Spontaneous thoughts and microstate SI modulation

To investigate how the observed SI-induced changes in microstates might reflect changes in the pattern of spontaneous thought, we conducted a partial least squares (PLS) analysis, a multivariate analysis that finds latent variables (LVs) which optimally link two sets of data by maximizing the covariance between the two. We found a significant LV (permuted p=0.012, 52% of covariance explained) for microstate *duration* (**Fig. 6a,b**). Changes in C and B microstate *duration* were positively correlated with thoughts about future plans (*planning*), and negatively associated with SI modulations of thoughts about *self* and thoughts about others. Microstates D and E showed the reversed patterns of associations (**Fig. 6c-e**).

**Figure 6.**
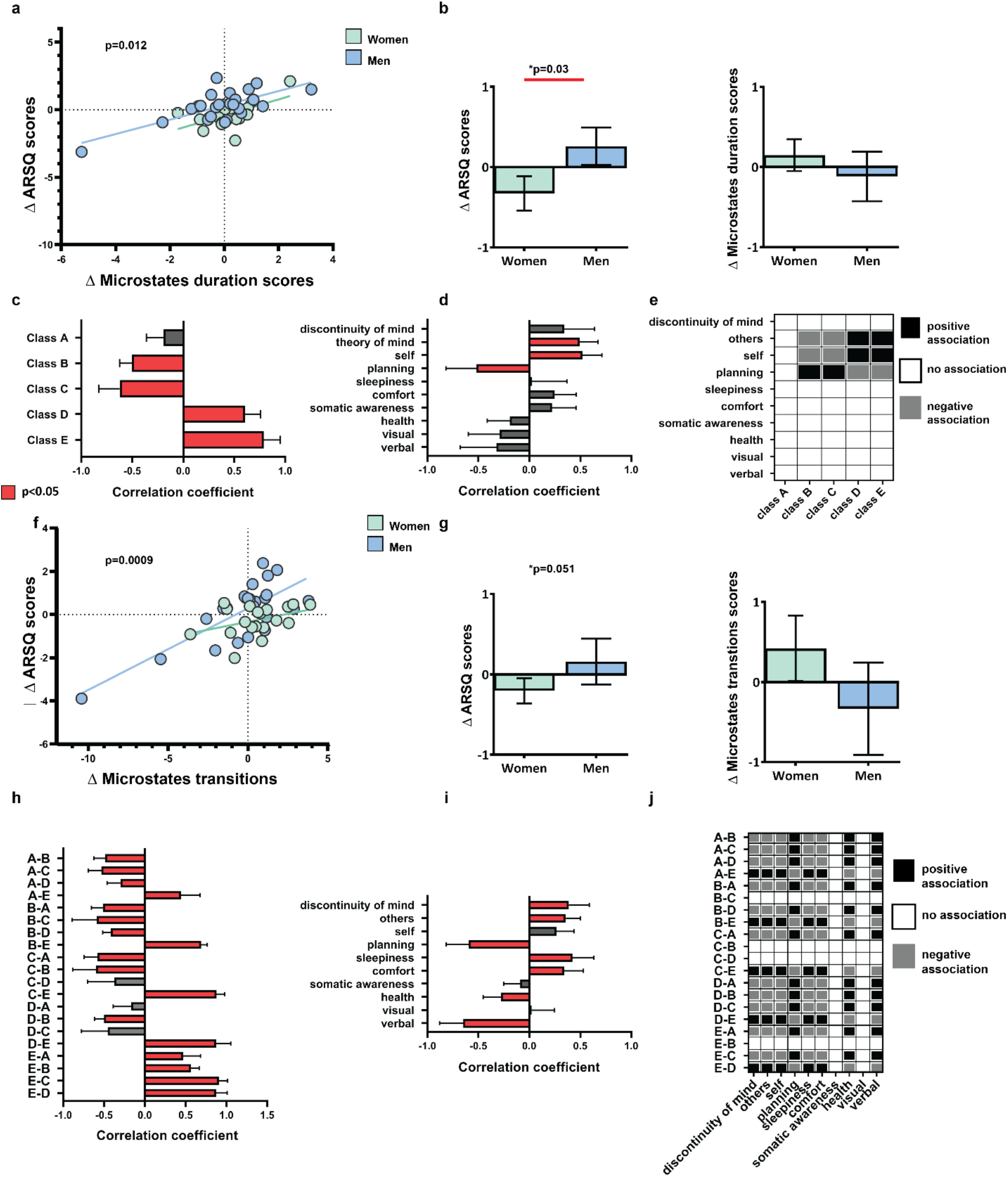
Association between microstate dynamics and spontaneous thoughts. (**a**) Correlation between individual-specific changes in ARSQ scores and changes in microstate duration. (**b**) Group differences in composite scores of ARSQ and microstate duration change. (**c**) Correlations between original and composite microstate duration scores (**d**) Correlations between original and composite ARSQ scores (**e**) Specific associations between microstate duration and spontaneous thoughts (**f**) Correlation between individual-specific changes in ARSQ scores and changes in microstate transitions. (**g**) Group differences in composite scores of ARSQ and microstate transitions change. (**h**) Correlations between original and composite microstate transition scores (**i**) Correlations between original and composite ARSQ scores (**k**) Specific associations between microstate transitions and spontaneous thoughts.

Results on *occurrence* modulations association with ARSQ change scores showed a significant LV (p=0.004, 54% explained covariance). Occurrence of microstates A and B was positively associated with *planning* and *verbal* thoughts. On the contrary, *occurrence* of microstate E was negatively associated with these aspects of spontaneous thoughts (**Fig. S1**).

We performed a separate PLS analysis using the SI dynamics of transition probabilities and SI dynamics of ARSQ scores in order to explore possible associations (**Fig. 6f-i**). We found a significant LV (p=0.0009, 53% explained covariance). For most transitions to A, B, C or D microstates we found a negative association with *discontinuity of mind*, *others*, *self*, *sleepiness*, *comfort* and a positive association with *planning*, *health* and *verbal* thoughts. From most microstates to E microstate, and for the E to D transitions, we found the reversed pattern of associations (**Fig 6j**). There were no associations found with *somatic awareness* and with *visual* thoughts.

These data indicate that the *duration* and *occurrence* of certain microstates could predict aspects of spontaneous thought content, whereas microstate *transition probability* could, in addition to content, also predict spontaneous thought dynamics, modality, and association with physiological states.

### Associations between microstate dynamics and salivary OXT following SI

To investigate if salivary OXT might point to a potential mechanism for how SI modulates microstate temporal features and transition probability, we tested if the dynamics of salivary oxytocin (ΔOXT) correlate with the dynamics of microstates duration, occurrence and transition probability (**Fig. 7**). There was no significant correlation between changes in OXT and either the *duration* or the *occurrence* of microstate classes (**Fig. 7c**). Similarly, in the whole group analysis there was no significant association between changes in OXT and transition probability (**Fig. 7b**). However, in women we found that ΔOXT negatively correlated with Δ A→C (Gamma r=-0.47, p=0.01), and positively correlated with Δ B→E (Gamma r=0.45, p=0.02). In men we found a positive association of Δ OXT with Δ D→B (Gamma r=0.32, p=0.03), as well as with Δ B→D (Gamma r=0.32, p=0.03).

**Figure 7.**
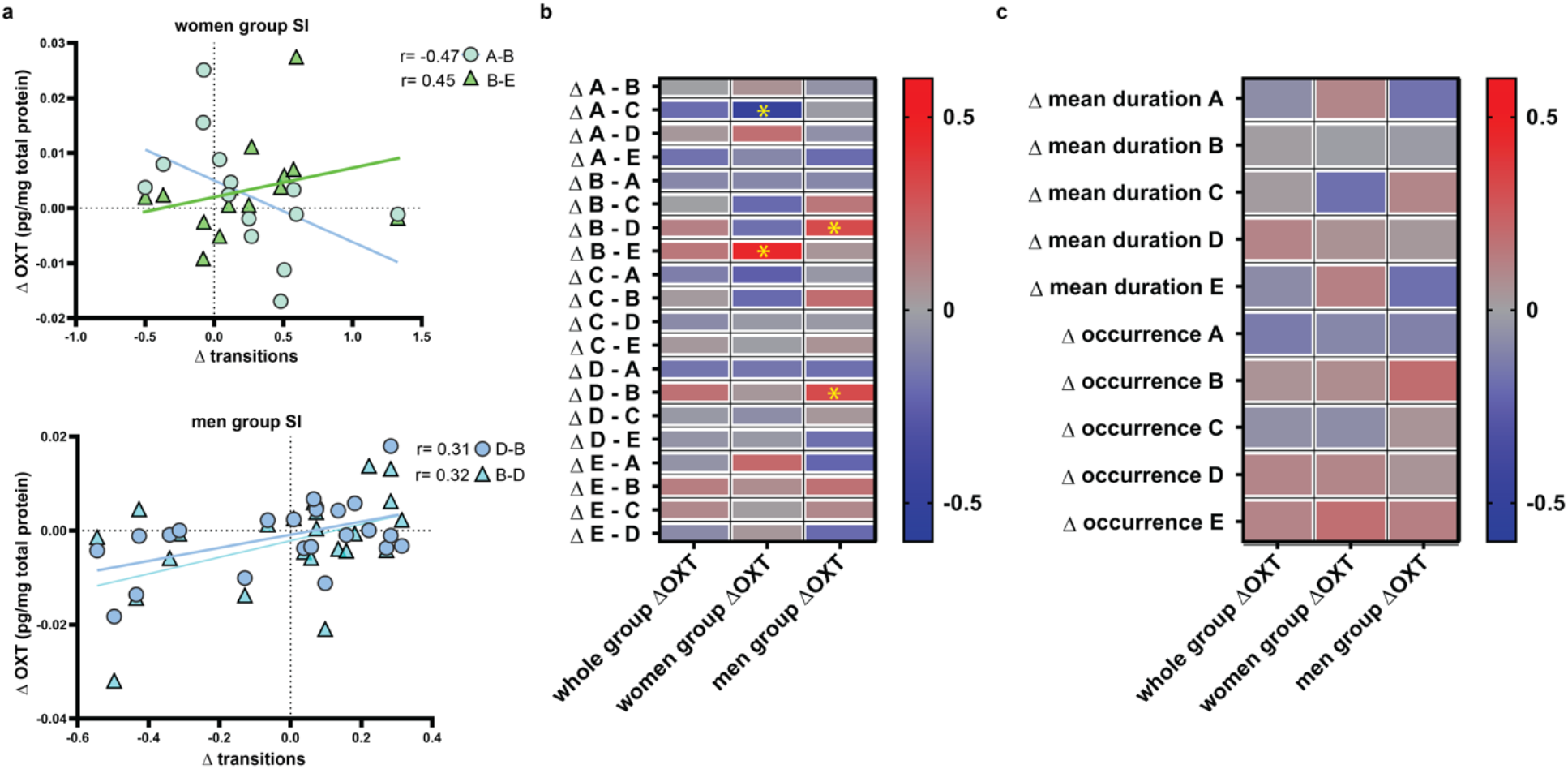
Association between changes in OXT and microstate temporal dynamics. (**a**) Example correlations between SI-specific changes in microstate transitions, and in salivary OXT (**b**) Summary of results for associations between changes in microstate transitions, and changes in salivary OXT following SI (**c**) Summary of results for associations between changes in microstate duration, occurrence, and changes in salivary oxytocin following SI.

### Associations between personality traits and temporal dynamics of EEG microstates

Responses to SI depend in part on personality traits (Papasteri, Sofonea et al. 2020). Similarly, personality traits impact spontaneous thought patterns (Diaz et al., 2014). Here, we investigated if SI modulations of spontaneous thoughts vary as a function of personality traits as measured by the NEO PI-R inventory. We found a significant LV (p=0.001, explaining 63% of covariance), revealing associations between personality traits, and SI modulation of ARSQ scores (**Fig. 2a-e**). *Neuroticism*, certain aspects of *openness* and *modesty* traits were positively associated with increases following SI of thoughts about self, of verbal thoughts and awareness of own physiological state during resting (**Fig. S2e**). *Extraversion* and *conscientiousness* traits were positively correlated with SI modulation of thoughts about planning and of visual imagery (**Fig. S2e**). These findings suggest that SI could be more beneficial for those with higher scores on extraversion and conscientiousness.

If changes in neural activity captured by changes in microstate properties serve as mechanisms for the effects of SI, we would expect to see associations between personality traits and microstate dynamics. Therefore, we conducted PLS analyses testing for associations between personality traits and microstates temporal dynamics. We found one LV (LV- p=0.007, 68% of covariance explained) showing that temporal dynamics of microstates following SI varied as a function of personality traits (**Fig. S3a**). Changes in microstates C, A, and B *duration* were positively related to *extraversion* and *conscientiousness* traits. The inverse pattern of association was present for microstate E. No significant LV was found explaining the *occurrence* changes as a function of personality traits. In summary, the significant changes in microstates *duration* following SI were more pronounced in those with higher scores in extraversion, conscientiousness and/or lower neuroticism, modesty and openness scores.

## Discussion

Imitation has been shown to have powerful effects on behavioural and emotional states in human and non-human primates^36,39^. So far, the search for a mechanism focused on changes in neural activity induces during imitation^37^. Here, we rely on previous findings linking mind-wandering to emotional states, to investigate if imitation might also act by changing the pattern of spontaneous thoughts. We find that imitation can indeed impact the dynamics, content and association with physiological states of mind-wandering episodes (**Fig. 1**), potentially by changing the pattern of correlated neuronal activity (**Fig. 4, 5)**. Further investigation is needed to determine if reported changes in spontaneous thoughts causally relate to changes in behavioural states and in salivary oxytocin.

### Association between spontaneous thoughts and behavioural state

We find that certain imitation-sensitive features of spontaneous thought dynamics and content positively associate with improvements in stress, well-being and closeness to the imitated person (**Fig. 1,2**). This finding adds the social implication to the reported effects of mind-wandering. Our findings on stress and well-being seem to contradict prior research. Previous studies report that negative mood and mind-wandering reinforce each other: spontaneous thoughts lead to sadness, which then leads to more mind-wandering ^7,10,11^. The apparent differences in our findings, namely the fact that mind-wandering can lead to improved behavioural states, could result from probing stress levels instead of affect valence, or from focusing on mind-wandering episodes around a generally agreeable task. The later possibility would support the notion that contexts have a substantial impact on spontaneous thoughts and on their ability to impact behaviour. The reverse sequence could also be true, where contexts impact behavioural states, which in turn modulate spontaneous thoughts. In this manuscript we do not establish a causal link, and further investigations will be needed to establish if such a link exists.

### Association with changes in OXT

The hormone OXT, known to play an important role in social bonding when acting centrally^40^, has been shown to increase following social imitation^36,38^. However, in previous work, we showed that while salivary OXT can predict well-being following SI, it does not associate with increased closeness for imitated partner^36^. This is consistent with our current findings where we report a negative association between salivary OXT and *discontinuity of mind*, a feature of mind-wandering that positively correlates with stress, and that decreases following imitation. One possibility is that increased OXT after SI acts on the dynamics of spontaneous thoughts to decreased stress levels. Also consistent with our previous report^36^, salivary OXT negatively associate with thoughts about planning, a correlate of increased social closeness following imitation. All results regarding salivary OXT should be interpreted with caution, as the levels of OXT in any peripheral fluid do not accurately reflect central actions of OXT^41,42^. Although the relationship between oxytocin and spontaneous thoughts has not been reported before, other hormones have been linked to mind-wandering. Changes in the level of the stress-related hormones cortisol and alpha-amylase have been associated with negative spontaneous thoughts^43^.

### Association with microstate dynamics

Our results show that changes in neural activity captured by microstate analysis could potentially explain changes in spontaneous thoughts and well-being following imitation. We find that the *duration* of microstate C, generally linked to activity in the DMN, increased after imitation (**Fig. 4**), and is positively associated with thoughts about planning, which also increase after imitation, and positively correlate with social closeness and well-being. Although we also observed changes in microstate D *occurrence* after imitation, they seemed to have less predictive potential for changes in spontaneous thought pattern and in behavioural states. However, the observed SI modulation of both microstate C and D are in line with previous findings on changes during relaxing behavioural states^32,33,34,35^. Changes in microstate transition probability following imitation were associated not only with changes in the content of spontaneous thought but also in their dynamics (**Fig. 6**), and in most cases were consistent with the effects of imitation on spontaneous thoughts. Additionally, we found a distinct pattern of association between SI modulation on microstate transitions and OXT (**Fig. 7**) in men compared to women. This supports previous data on the sexually dimorphic action of OXT, and on the role of microstate dynamics underlying different patterns of information processing^44,45^. In agreement with previous finding that salivary OXT increases in women but not men following SI^36^, we further show that only in women the change in OXT is associated with slower transitions from microstate A (generally linked to activity in language/auditory cortices) to microstate C (associated to posterior DMN activations)^26,31^.

Could ongoing, stimulus- and task-independent brain activity impact spontaneous thoughts and possibly subsequent behavioural states? The impact of ongoing neuronal activity on task-related performances has been studied extensively in both animal models and humans^46,47^, yet the association between ongoing activity patterns and spontaneous thoughts has been more difficult to establish, due to the fleeting nature and unpredictability of spontaneous thoughts. In addition to our current findings linking ongoing patterns of microstate activity with dynamics and content of spontaneous thoughts, previous work showed that patients suffering from neuropsychiatric disorders with abnormal mind-wandering patterns have significant differences in microstate dynamics^48–54^.

Taken together, our findings highlight the effect of social imitation on spontaneous thoughts, and the potential neural and biochemical substrates involved. Our findings should fuel research into potential therapies using social context-dependent changes in spontaneous thoughts.

## Methods

### Ethical considerations

All methods and experiments have been approved by The Ethics Committee of National University for Theatre and Film I.L Caragiale Bucharest, and followed the guidelines of the Declaration of Helsinki. All participants provided written informed consent for their participation. The instructor and subject shown in **Figure 1a**, both gave their written consent to have their faces shown in the manuscript.

### Experimental paradigm

This study investigates social imitation (SI) task modulations of resting-state EEG, OXT and behavioural states. Data from each participant was assessed across the different tasks (SI vs CTRL) on two separate days in a counterbalanced, randomized fashion, with at least one week delay between the two experimental days.

As depicted in Figure 1, during the SI task participants were asked to imitate in real time the stereotyped, geometrical arm movements produced by an instructor (e.g. draw a circle in the air). The SI task lasted for 3 minutes during which the instructor provided positive verbal cues (e.g. “very well”) and positive non-verbal cues (e.g. smile, nodding yes). During CTRL task no social interaction was available and participants were asked to imitate the geometrical movements displayed on a screen (3 minutes).

### Participants and data collection

Participants in the study were recruited through advertisement within the University of Bucharest, University of Theatre and Film and the CINETic’s Research Centre website https://cinetic.arts.ro/en/met-2/. The exclusion criteria included neurologic and psychiatric symptoms. Of the 65 subjects that were initially recruited, several (N=7) were excluded from the analysis due to movement contamination of the EEG data and several (N=15) dropped out of the experiment before the collection of the second day of the experiment. The remaining dataset thus included 43 participants (24 men, 19 women), mean age=25.7 (age range: 20-42), s.d.=5.1. (Women mean age=25.57, s.d.=5.3; Men mean age=25.9, s.d.=5.1). The sample group included in the correlation analyses between OXT or personality traits and EEG temporal dynamics was composed of 37 participants (18 men and 19 women). Six participants were excluded from these analyses due to missing OXT and NEO-PI-R data.

EEG data were acquired in a dimly light room using a 128-channel ANT Neuro waveguard™ system (https://www.ant-neuro.com/) sampled online at 1kHZ with a Cz reference. Participants sat in a comfortable, upright position and were instructed to stay awake, as calm as possible, to keep their eyes closed and to relax for five minutes without falling asleep.

Salivary oxytocin (OXT) were collected using specially designed tubes (Salivette©, Sarsted). Participants were instructed to saturate the saliva swab by holding for at least 2 minutes in the mouth. The tubes were then centrifuged at 1000 x g, at 4° C, for 25 minutes and the samples were aliquoted in 1,5 ml Eppendorf vials and stored at −80° C prior to analysis. Oxytocin was measured by radioimmunoassay (RIA) at RIAgnosis, Munich, Germany, while total proteins were measured at National Institute of Endocrinology “C. I. Parhon”, Bucharest, Romania. Salivary total protein was used to normalize the concentration of salivary oxytocin levels, since its concentration can vary significantly with saliva viscosity.

Self-reported data were collected in the same room as the EEG data and consisted of several pen-paper tests. The inclusion of others in the self (IOS)^55^ a single item measure of closeness composed of two Venn-like circles varying in their degree of overlap. Participants were instructed to select the diagram reflecting the relationship with the instructor that guided the participants throughout the social interaction task. Two visual analog scales (VAS) assessed the self-reported level of stress and well-being using a10-cm unmarked scale ranging from 0 “no stress”/”worst unimaginable well-being” up to 100 “most stress ever”/”perfect well-being”.

The Neo Personality Inventory-Revised (NEO PI-R) is a 240 item personality inventory with cross-culturally established psychometric properties and validity^56^ that assesses the Big Five Model domains of Neuroticism, Extraversion, Openness to Experience, Agreeableness and Conscientiousness with the 6 factors for each domain. Participants responded on a 5-point Likert scale ranging from 0 (strongly disagree) to 4 (strongly agree).

The Amsterdam Resting-State Questionnaire 2.0 is a self-report questionnaire that quantifies mind wandering along ten model: Discontinuity of Mind, Theory of Mind, Self, Planning, Sleepiness, Comfort, Somatic Awareness, Health Concern, Visual Thought, Verbal Thought^9^. Participants had to responded to a total of 30 items using a 5-point Likert Scale from “Completely Disagree” to “Completely Agree”.

### EEG data processing

The EEG datasets were band-pass filtered offline between 1 and 40 Hz with an additional notch at 50 Hz. EEG periods of movement contamination or other artifacts were marked and excluded from the analyses. In order to remove the oculomotor artifacts such as saccades and eye blinks as well as the cardiac artefacts (ECG), we applied the Infomax-based Independent Component Analysis (ICA)^57^. Bad or noisy electrodes were interpolated using a 3-D spherical spline^58^, and were recomputed to the common average-reference. The data were then down-sampled to 125 Hz for further analysis.

The local maxima of the Global Field Power (GFP) show an optimal signal to noise-ratio in the EEG^58,59^. The EEG signal was extracted at the corresponding time frame of GFP peaks and only the time points of GFP peaks were submitted to a modified k-means cluster analysis^59,60^ in order to identify the most representative classes of stable topographies.

The k-means clustering was performed in two steps: first, at the individual level, and, in a second step, at the group level by clustering all individual dominant topographies with varying number of clusters. In order to determine the optimal number of clusters at the individual and the group level, we used the criteria implemented in Cartool (a free academic software developed by Denis Brunet; <u>cartoolcommunity.unige.ch</u>), based on seven maximally independent criteria: Davies and Bouldin, Gamma, Silhouette, Dunn Robust, Point-Biserial, Krzanowski-Lai Index, and Cross-Validation ^31,59,61^.

In the first part of the microstate analysis, only GFP peaks were submitted to the k-means clustering. However, in the second part of the analysis, during the fitting process of the microstates, the entire EEG of participants was used, excluding only the marked artifact epochs. A temporal smoothing (window half-size 3 (24 ms), Besag factor of 10 and a rejection of small time frames (when <3, i.e. 24 ms) was applied^59^. Subsequently, in order to quantify the temporal parameters of microstates, every time point of the individual data was assigned to the microstate cluster with which it correlated best^60^. A 0.5 correlation coefficient threshold was used in order to exclude transient periods of noise in the data. These periods were not labelled and were excluded from the analysis.

This fitting process enabled the determination of the duration and the occurrence of each microstate in each subject. The *duration* represents the average amount of time (in ms) that a given microstate map was present without interruptions, i.e. the duration during which the subject remained in a certain state. The duration is one of the most commonly used parameters of the temporal structure of microstates and has repeatedly been shown to be associated with different vigilance conditions and symptoms of neuropsychiatric disorders^62^. The mean *occurrence* of a microstate is independent of the duration. It indicates the rate at which a given microstate occurred, i.e. how many times per second the brain enters a certain state.

In addition to these two temporal parameters for each microstate, we also analysed the between microstate transition dynamics: for each subject and transition pair we computed the number of transitions and normalized them by all between-class transitions^45,63^.

The free academic software Cartool (<u>cartoolcommunity.unige.ch</u>) was used for the EEG data processing and microstate analysis^61^.

### Statistical analyses

In order to assess the specific modulations of spontaneous thoughts after the SI we utilized a three-way repeated measures ANOVA with three factors: the ARSQ factors (10), time (PRE-POST), and task (SI or CTRL). The averaged ARSQ scores across the different experimental conditions (PRE-SI, POST-SI, PRE-CTRL, POST-CTRL) were averaged for each individual and correlation analyses to investigate associations with behavioural states (stress, well-being and IOS) and salivary OXT. We used Gamma non-parametric correlation analyses which can inform us on the strength of their association disregarding possible outliers and tied ranks in the data.

Similarly, for both microstate temporal parameters (duration and occurrence) and transition probabilities for each directional transition pair (20). Separate three-way repeated measures ANOVA’s with time (PRE-POST), classes of microstates (5)/transition pairs (20), and task (SI or CTRL) as within factors was performed. We performed non-parametrical Wilcoxon paired test to further assess main drivers of significant main effects. Additionally, to assess the degree of gender differences of these effects we performed separate statistical analyses on men and women subgroups.

In order to identify robust patterns of correlations between the behavioural measures and the temporal dynamics of the EEG microstates, we used a multivariate approach called partial least squares (PLS^)64^. PLS is a multivariate data-driven statistical technique that maximizes the covariance between two matrices by deriving *latent variables* (LVs), which are optimal linear combinations of the original matrices^65^. PLS is a powerful technique for relating two sets of data (e.g., neuroimaging and behavioral data), even if these data show autocorrelation or multicollinearity^64^.

Each LV is characterized by a distinct EEG microstate pattern (called EEG *loadings*) and a distinct behavioral profile (called behavioral *loadings*). By linearly projecting the EEG and behavioral measures of each participant onto their respective loadings, we obtained individual-specific EEG microstates and behavioral *composite scores* for each LV. PLS seeks to find loadings that maximize across-participant covariance between the EEG microstates parameters and behavioral composite scores. The number of significant LVs was determined by a permutation test (1000 permutations). The p-values (from the permutation) for the first five LVs were corrected for multiple comparisons using a FDR of q < 0.05. To interpret the LVs, we computed Pearson’s correlations between the original EEG data and EEG composite scores, as well as between the original behavioral measures and behavioral composite scores for each LV. A large positive (or negative) correlation for a particular behavioral measure for a given LV indicates greater importance of the behavioral measure for the LV. Similarly, a large positive (or negative) correlation for a particular EEG microstate parameter for a given LV indicates greater importance of the EEG microstate parameter for the LV. To estimate confidence intervals for these correlations, we applied a bootstrapping procedure that generated 500 samples from subjects’ data. Z-scores were calculated by dividing each correlation coefficient by its bootstrap-estimated standard deviation. The z-scores were converted to p-values and FDR-corrected (p < 0.05)^66^.

## AUTHOR CONTRIBUTIONS

All authors contributed to the design of the experiment and interpretation of results. MIT (first author) analyzed all data, in collaboration with CP and VK. MIT, AS, RB conducted the experiments and data collection. MIT and IC wrote the manuscript, with feedback from all authors.

## ACKNOWLEDGEMENT

We thank Prof. Nicolae Mandea, Prof. Liviu Lucaci and Prof. Carmen Croitoru for their administrative support, Dr. Robert C. Froemke and Dr. Justin S. Riceberg for consultation, Nicoleta Puşcaşu and Doina Strat for their technical support. The Cartool software (cartoolcommunity.unige.ch) has been programmed by Denis Brunet, from the Functional Brain Mapping Laboratory (FBMLab), Geneva, Switzerland, and is supported by the Center for Biomedical Imaging (CIBM) of Geneva and Lausanne.

