## Supplemental material for "Spontaneous thought and microstate activity modulation by social imitation"

**Supplementary figures**

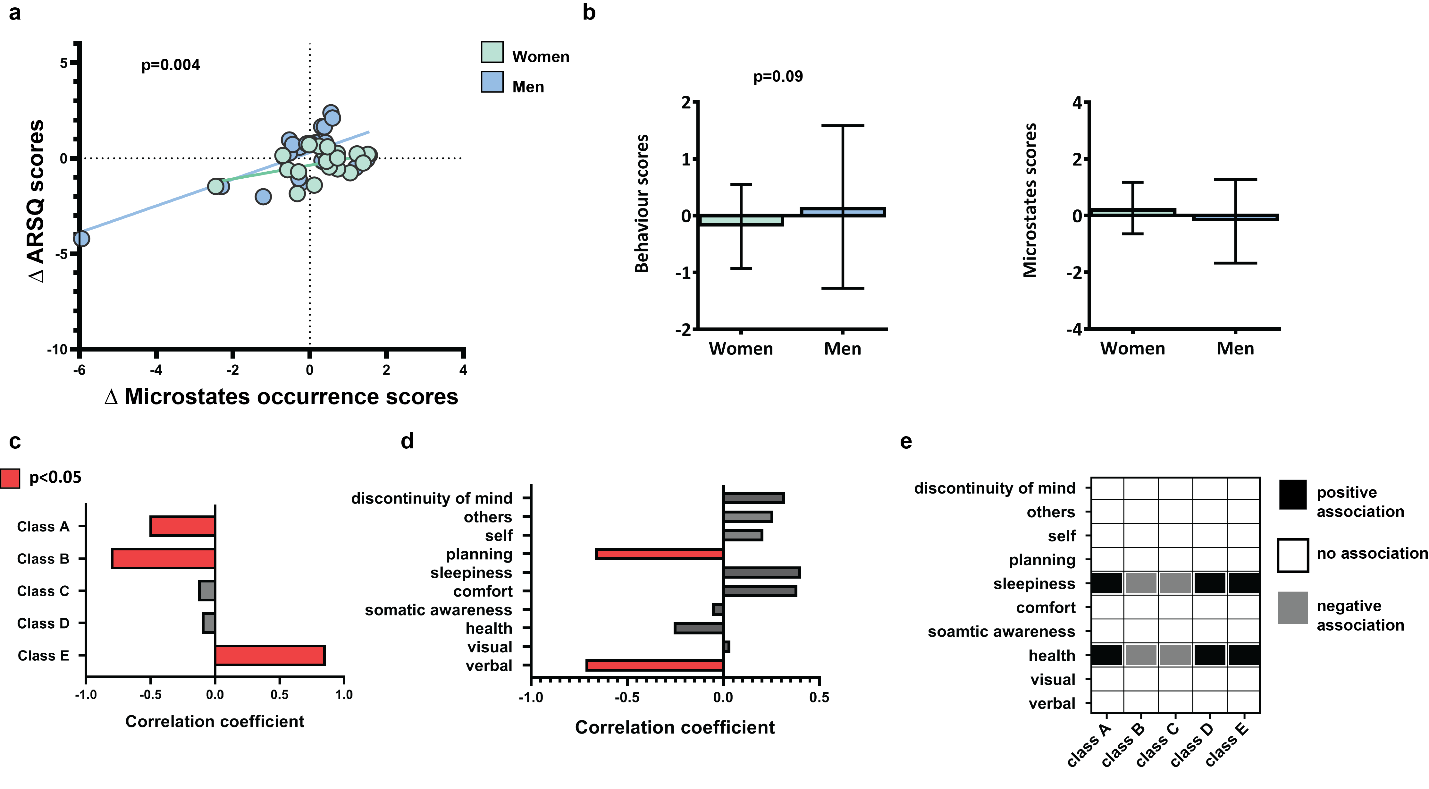

**Figure S1. Association between microstate occurrence and spontaneous thoughts.**

(**a**) Correlation between individual-specific changes in ARSQ scores and changes in microstate occurrence. (**b**) Group differences in composite scores of ARSQ and microstate occurrence change. (**c**) Correlations between original and composite microstate occurrence scores. (**d**) Correlations between original and composite ARSQ scores. (**e**) Specific associations between microstate occurrence and spontaneous thoughts.

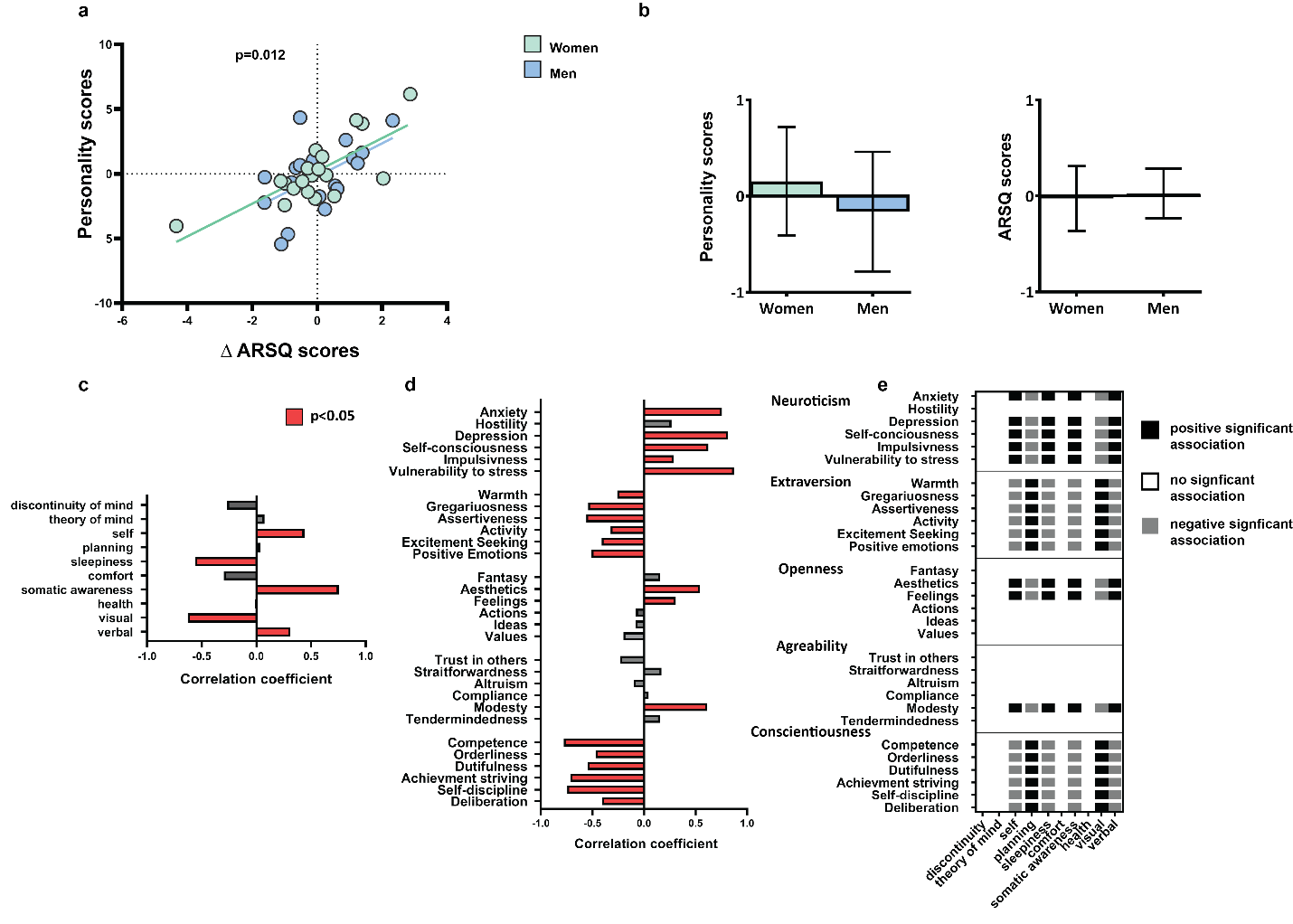

**Figure S2. Association between personality traits and spontaneous thoughts.**

(**a**) Correlation between individual-specific changes in ARSQ scores and personality traits. (**b**) Group differences in composite scores of personality traits and ARSQ change scores.

(**c**) Correlations between original and composite ARSQ scores. (**d**) Correlations between original and composite personality traits. (**e**) Specific associations between personality traits and spontaneous thoughts.

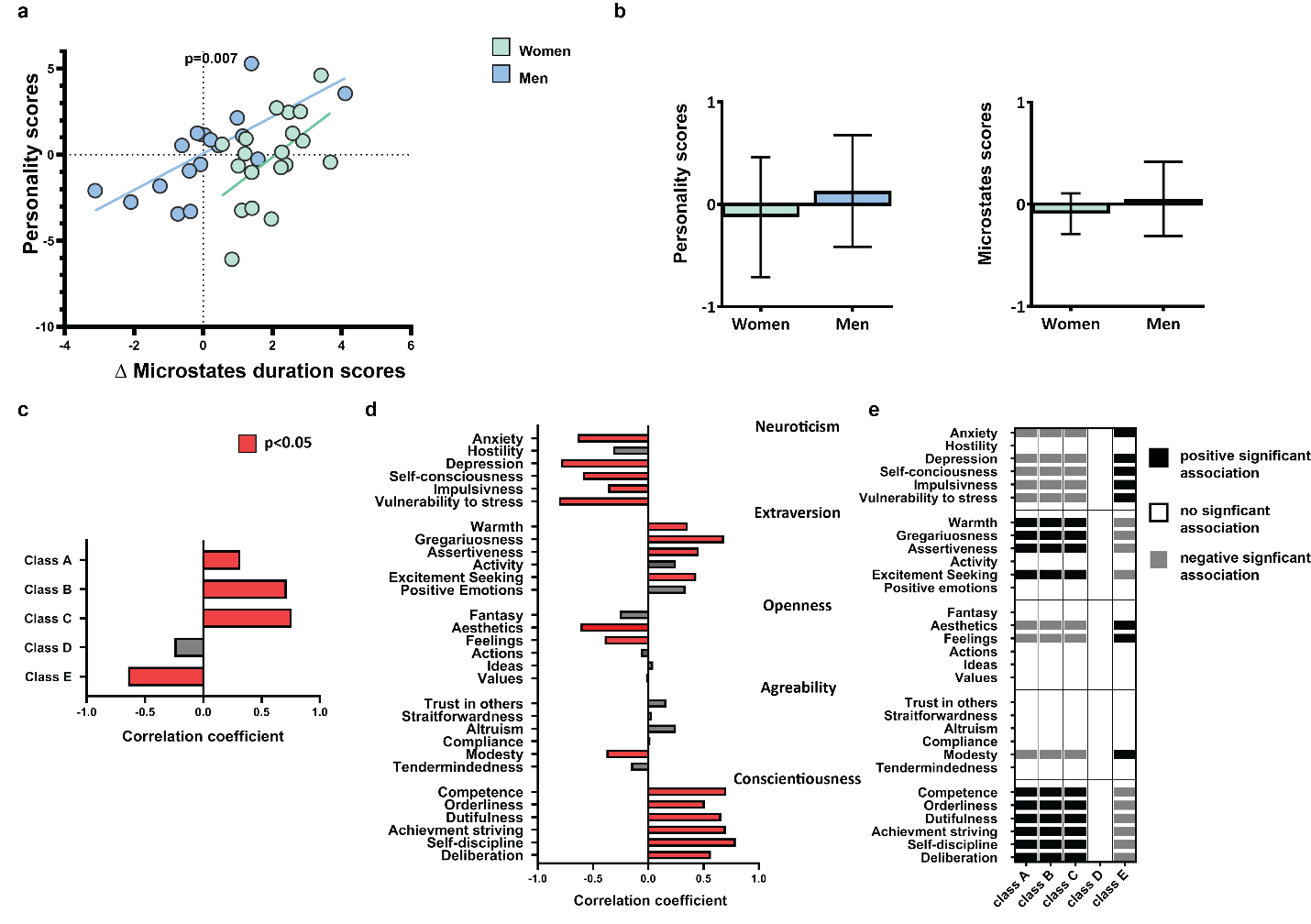

**Figure S3. Association between personality traits and microstate duration.**

(**a**) Correlation between individual-specific changes in microstate duration and personality traits. (**b**) Group differences in composite scores of personality traits and microstate duration change scores. (**c**) Correlations between original and composite microstate duration scores. (**d**) Correlations between original and composite personality traits. (**e**) Specific associations between personality traits and spontaneous thoughts.
